# PCNA Antagonizes Cohesin-dependent Roles in Genomic Stability

**DOI:** 10.1101/2020.06.19.161489

**Authors:** Caitlin M. Zuilkoski, Robert V. Skibbens

## Abstract

PCNA sliding clamp binds factors through which histone deposition, chromatin remodeling, and DNA repair are coupled to DNA replication. PCNA also directly binds Eco1/Ctf7 acetyltransferase, which in turn activates cohesins and establishes cohesion between nascent sister chromatids. While increased recruitment thus explains the mechanism through which elevated levels of chromatin-bound PCNA rescue *eco1* mutant cell growth, the mechanism through which PCNA instead worsens cohesin mutant cell growth remains unknown. Possibilities include that elevated levels of long-lived chromatin-bound PCNA reduce either cohesin deposition onto DNA or cohesin acetylation. Instead, our results reveal that PCNA increases the levels of both chromatin-bound cohesin and cohesin acetylation. Beyond sister chromatid cohesion, PCNA also plays a critical role genomic stability such that high levels of chromatin-bound PCNA destabilize the DNA replication fork. At a relatively modest increase of chromatin-bound PCNA, however, fork stability and progression appear normal in wildtype cells. Our results reveal that even this moderate increase of PCNA sensitizes cohesin mutant cells to DNA damaging agents and in a process that involves the DNA damage response kinase Mec1(ATR), but not Tel1(ATM). These and other findings document that PCNA mis-regulation results in genome instabilities that normally are resolved by cohesin and suggest that elevating levels of chromatin-bound PCNA may target cohesinopathic cells linked to cancer.

## INTRODUCTION

During S phase, the cellular genome duplicates and each sister chromatid becomes tethered together to ensure proper inheritance of the genome during mitosis. Sister chromatid tethering is maintained by cohesin, a complex comprised of Smc1, Smc3, and Mcd1/Scc1/RAD21 along with auxiliary subunits Pds5, Scc3/Irr1, Rad61/WAPL and, in vertebrate cells, Sororin [1–7]. Cohesin deposition onto DNA requires the cohesin loader, comprised of Scc2,4, which functions through a large part of the cell cycle but is essential during S phase for cohesins to participate in sister chromatin tethering [8,9]. Deposition onto chromatin, however, is not sufficient for cohesion. Eco1/Ctf7 (herein Eco1) is an acetyltransferase that converts chromatin-bound cohesins, through acetylation of Smc3 cohesin subunit, to a tethering competent state [3,10–13]. Early studies coupled this process of cohesion establishment to the DNA replication factor PCNA [10]. PCNA directly binds and recruits Eco1 to the DNA replication fork [14–16], suggesting that establishment is coordinated with numerous processes (histone deposition, chromatin remodeling, DNA repair and translesion synthesis) directed by PCNA and regulated through PCNA post-translational modifications [17–20]. Elevated recruitment to the DNA replication fork thus provides a plausible explanation for PCNA-dependent suppression of *eco1* mutant cell growth defects [10,15]. The repertoire of DNA replication factors (Chl1, Ctf4, Ctf18, MCM2-7, for example) implicated in cohesion establishment regulation has grown substantially [15,21–35], highlighting the fundamental and highly conserved nature through which cohesion establishment is obligatorily coordinated with DNA replication.

Further complicating the relationship between PCNA, Eco1 and cohesins are the additional roles played beyond sister chromatid tethering [1,5,10,36–47]. For instance, cohesin functions in both DNA replication restart during S phase and high fidelity DNA repair after S phase [48–50]. Cohesin is recruited to stalled DNA replication forks during S phase, as well as to rDNA and telomeric regions that experience prescribed pauses during replication [51–53]. Cohesin is similarly recruited to sites of DNA damage to promote high fidelity repair by ensuring proximity to the undamaged sister template [49,54,55]. Intriguingly, once recruited to stalled replication forks and sites of DNA damage, cohesin must ultimately dissociate from DNA to promote replication fork restart [56]. Thus, cohesin and PCNA appear to play highly coordinated roles in DNA metabolism.

PCNA impact on cohesin complexes, however, is quite complex. On the one hand, deletion of the PCNA-dissociation factor *ELG1* (*elg1Δ*), or simply overexpression of PCNA (PCNA^OE^), result in long-lived retention and increased levels of chromatin-bound PCNA which rescue both *eco1* and *pds5* mutant cell cohesion defects and viability [10,15,31–33,57–62]. On the other hand, *elg1Δ* and PCNA^OE^ each exacerbate the growth defects exhibited by cohesin (*mcd1, smc1* and *smc3*) mutant cells [15,31,32,34]. The mechanism through which elevated levels of chromatin-bound PCNA antagonize cohesin mutant cell growth remains unknown, highlighting a critical deficit in current descriptions of cohesin regulation. In this current study, we confirm the adverse impact that PCNA^OE^ exerts on cohesin mutant strains even while having no overt effect in wildtype cells. We then test multiple mechanisms through which the long-lived retention increased level of chromatin-bound PCNA exacerbates cohesin mutant cells, including competition for DNA access, changes in cohesin acetylation states and increased genomic instability.

## MATERIALS AND METHODS

### Yeast Strains and Bacterial Plasmids

All reagents, yeast strains, and bacterial plasmids are listed in Tables Supplemental 1, 2, and 3, respectively. All yeast strains used in this study were performed in a W303 background strain unless otherwise noted (Supplemental Table 2). V5 tagged Smc3 strains were created using EU3430-9A, generously given by Drs. Vincent Guacci and Douglas Koshland. Primers used to amplify the SMC3:V5:HIS region, primers are (forward primer) 5’-TTAACGCGGTTGATTTCTACTTTCCAAAAGGTTTCTGAAAA-3’and (reverse primer) 5’-TAGCTCTGATTCTGACTCTAACTCCAGTTCGGACTCCGTATCGGATTCCAGTTCAGATTC-3’. The resulting PCR product was transformed into a wildtype W303 background strain (YCZ044) to produce YCZ249. YCZ249 was mated with YCZ262 and a genetic cross was used to obtain YCZ280. YCZ280 was mated with YBS2012 to obtain YCZ407, YCZ408, YCZ425, and YCZ428 (see Supplemental Table 2). A CEN vector plasmid or *2µ POL30* plasmid were transformed into YCZ147 and YCZ144 to obtain the resulting yeast strains: YCZ477, YCZ465, YCZ474, and YCZ225. YCZ559, YCZ561, YCZ563, and YCZ565 were created by transforming either a CEN vector plasmid or *2µ POL30* plasmid into K5824 and K6013, respectively, generously given by Dr. Vincent Guacci. A CEN vector plasmid or *2µ POL30* plasmid were transformed into YMM433 and YMM435 to obtain the resulting yeast strains (YCZ530, YCZ532, YCZ534, YCZ536). A CEN vector plasmid or *2µ POL30* plasmid were transformed into YDM884 and DLY285 to obtain the resulting yeast strains (YCZ567 through YCZ578). A pGADT7 vector plasmid or a *2µ ADH AD:HA:POL30* plasmid was transformed into yeast strains K699, K5824, and K6013 to generate resulting yeast strains (YCZ662 through YCZ673). *2µ POL30* plasmid was transformed into yeast strains K5824 and K6013 to generate resulting yeast strains (YCZ559, YCZ561, YCZ563, YCZ565). Primers (forward primer) 5’-AGAGAAGGTTTTCCAATGAAAAGGCACGTGTCTTTATCTGATATATTGACAGGAAATAAGCGGATCCCCGGGTTAATTAA-3’and (reverse primer) 5’-ATTTCCCCGCACTACCATGCTATATTTATTATACATACGTGTTCCTGTAACGATGCACGCGAATTCGAGCTCGTTTAAAC-3’ were used to generate strains harboring *ELG1::KAN* as previously described [63]. The resulting PCR product was transformed into K5824 to obtain the yeast strains YCZ706 and YCZ707. YCZ143 was mated to K6013 to obtain YCZ702 and a genetic cross was used to obtain YCZ728 and YCZ729. CZ421 was mated to CZ778, generously given by Dr. Gregory Lang, to obtain CZ799 and a genetic cross was used to determine the genetic interaction between *msh3Δ, elg1Δ*, and *mcd1-1* alleles. Strains and plasmids are available upon request.

### Dilution plating

Cells were grown at 23° overnight in either YPD or selection media, normalized by OD600, and then plated in 10-fold serial dilutions at the indicated temperatures. Cells grown on YPD medium or selective medium supplemented with methyl methanesulfonate (Sigma) or hydroxyurea (Sigma) were treated in a similar manner.

### Chromatin binding assay

Log phase yeast strains were grown to 0.6 OD600 and synchronized in G1 by exposing cells to fresh media supplemented with alpha factor at the permissive temperature of 23°C for three hours. The resulting G1 synchronized cells were washed, cultures split in half and maintained for three hours at 34°C or 37°C for 3 hours. Nocodazole arrested cells were harvested to assess Smc3 chromatin binding levels, based on modifications of previously described protocols [64]. Cell cultures were normalized to OD600 between 0.3–0.6, pelleted and washed with 1.2 M Sorbitol. Cells were resuspended in CB1 buffer (50 mM Sodium citrate, 1.2 M Sorbitol, 40 mM EDTA, pH 7.4) and spheroblasted. Spheroblasted cells were washed, resuspended in 1.2 M Sorbitol, and frozen in liquid nitrogen. Cells were thawed on ice and supplemented with Lysis buffer (500 mM Lithium Acetate, 20 mM MgSO4, 200 mM HEPES, pH 7.9), protease inhibitor cocktail (AEBSF, 1,10-Phenanthroline, Pepstatin A, E-64) (Sigma), and TritonX-100. Lysates were centrifuged at 12,000g for 15 minutes and soluble and containing chromatin bound fractions collected and denatured using 2X Laemelli buffer. Whole cell extracts, supernatant, and pellet were resolved by SDS-PAGE and analyzed by Western blot using anti-V5 (1:40,000) (Invitrogen) with goat anti-mouse HRP (1:40,000) (Bio-Rad), anti-PGK (1:20,000) (Novex) with goat anti-mouse HRP (1:40,000) (Bio-Rad), or anit-H2B (1:80,000) (Abcam) with goat anti-rabbit HRP (1:40,000) (Bio-Rad) and ECL prime (GE Healthcare) for visualization.

### Smc3 acetylation assay

Log phase yeast strains were grown to 0.6 OD600 and synchronized in G1 by exposing cells to fresh media supplemented with alpha factor at the permissive temperature of 23°C for three hours. The resulting G1 synchronized cells were washed, cultures split in half and maintained for three hours at 34°C or 37°C for 3 hours. Nocodazole arrested cells were harvested to assess Smc3 acetylation, with additional modifications as previously described [64]. Cell cultures were normalized to an OD600 between 0.3–0.6. Cells were washed in sterile water, then resuspended in sterile water prior to freezing at −80°C. Frozen pellets were extracted by the addition of IPH50 buffer (50mM Tris pH 7.8, 150mM NaCl, 5mM EDTA, 0.5% IGEPAL 630 (Sigma), 10mM Sodium Butyrate, 1mM DTT) and glass beads prior to bead beating (BioSpec). Cell lysates were supplemented with IPH50 buffer and protease inhibitor cocktail (AEBSF, 1,10-Phenanthroline, Pepstatin A, E-64) (Sigma), centrifuged at 15,000rpm for 20 minutes, and the pellet washed with sterile water before centrifugation at 15,000rpm for 10 minutes. The resulting chromatin fraction was supplemented with SBIIA buffer (0.5M Tris pH 9.4, 6% Sodium Dodecyl Sulfate before a 10-minute incubation at 50°C. SBII buffer (50% glycerol supplemented with bromophenol blue) and 1M DTT buffer was added to the cell lysates followed by a 5-minute incubation at 65°C. Whole cell protein samples were resolved by SDS-PAGE electrophoresis and analyzed by Western blot using anti-V5 (1:40,000) (Invitrogen) with goat anti-mouse HRP (1:40,000), anti-PGK (1:20,000) (Novex) with goat anti-mouse HRP (1:40,000) (Bio-Rad) or by anti-Smc3 K112/K113 Acetylation (1:1,000, gift from Dr. Katsuhiko Shirahige) in combination with goat anti-mouse HRP (1:10,000) (Bio-Rad) and ECL prime (GE Healthcare) for visualization.

## RESULTS

### Elevated levels of chromatin-bound PCNA (via *elg1Δ*) promote cohesin binding to DNA in *mcd1* mutant cells

PCNA^OE^ (expressed from a 2*µ* vector) and *elg1Δ*, both of which result in persistent and increased levels of chromatin-bound PCNA, adversely affect all core cohesin mutant strains tested to date [15,31,32,34,60,62]. Given that cohesin deposition onto chromatin, and cycles of PCNA binding/release, are normally coordinated, it became important to test whether elevated levels of chromatin-bound PCNA reduce cohesin deposition onto DNA - either through direct competition, reduction of factors that promote cohesin deposition (Scc2,4, RSC or Chl1), or increase in factors (Rad61/WAPL) that dissociate cohesin from chromatin [7,8,22,29,30,62,65– 69]. We first independently verified the antagonistic effect that *elg1Δ* imparts on cells harboring *mcd1-1* mutations and indeed found severely exacerbated growth defects (Figure 1A). Next, we validated the efficiency of the Triton X-100-based fractionation procedure [3,44]. We obtained clear separation of soluble components (using the cytoplasmic factor Phosphoglycerokinase - PGK) from those of chromatin-bound components (Histone 2B) (65). Finally, we optimized Western blot procedures to obtain a linear range of band intensities from which to quantify levels of fractionated chromatin-bound proteins (Figure 1B). To test whether elevated levels of chromatin-bound PCNA reduce cohesin binding to DNA, wildtype, *elg1Δ* and *mcd1-1* single mutant cells, and *mcd1-1 elg1Δ* double mutant cells, all of which express Smc3-3V5 as the sole source of Smc3 protein, were synchronized in G1 and released at either 34°C or 37°C (non-permissive for *mcd1-1 elg1Δ* double mutant cells and *mcd1-1* single mutant cells, respectively) in fresh medium containing nocodazole. Cell cycle synchronizations were monitored by flow cytometry (Figure 1C). The resulting pre-anaphase cells were lysed and fractionated prior to assessing for changes in chromatin-bound Smc3, a core component of the cohesin complex [2,70]. As expected, wildtype and *elg1Δ* single mutant cells display similar levels of chromatin bound Smc3, compared to H2B (Figure 1D and E). The results further reveal that *mcd1-1* single mutant cells display similar levels of chromatin-bound Smc3 compared to wildtype cells, even at the non-permissive temperature of 37° (Figure 1D and E). Thus, Mcd1 inactivation, which results in cell inviability, cohesion and condensation defects [1,2], does so in the absence of cohesin dissociation from DNA (Figure 1D and E). Surprisingly, *mcd1-1 elg1Δ* double mutant cells display increased levels of chromatin bound cohesin at both non-permissive temperatures of 34° and 37°, compared to that of either wildtype, *elg1Δ* or *mcd1-1* single mutant cells (Figure 1D and 1E). These results that chromatin-bound levels of Smc3 are not reduced in *mcd1-1 elg1Δ* double mutant cells, but instead are increased, reveal that excess chromatin-bound PCNA does not adversely impact cohesin loading onto DNA and negate both direct competition models and recruitment of cohesin releasing factors. The mechanism through which cohesin binding to chromatin is enhanced by *elg1Δ* in *mcd1-1* mutant cells remains unknown.

**Figure 1:**
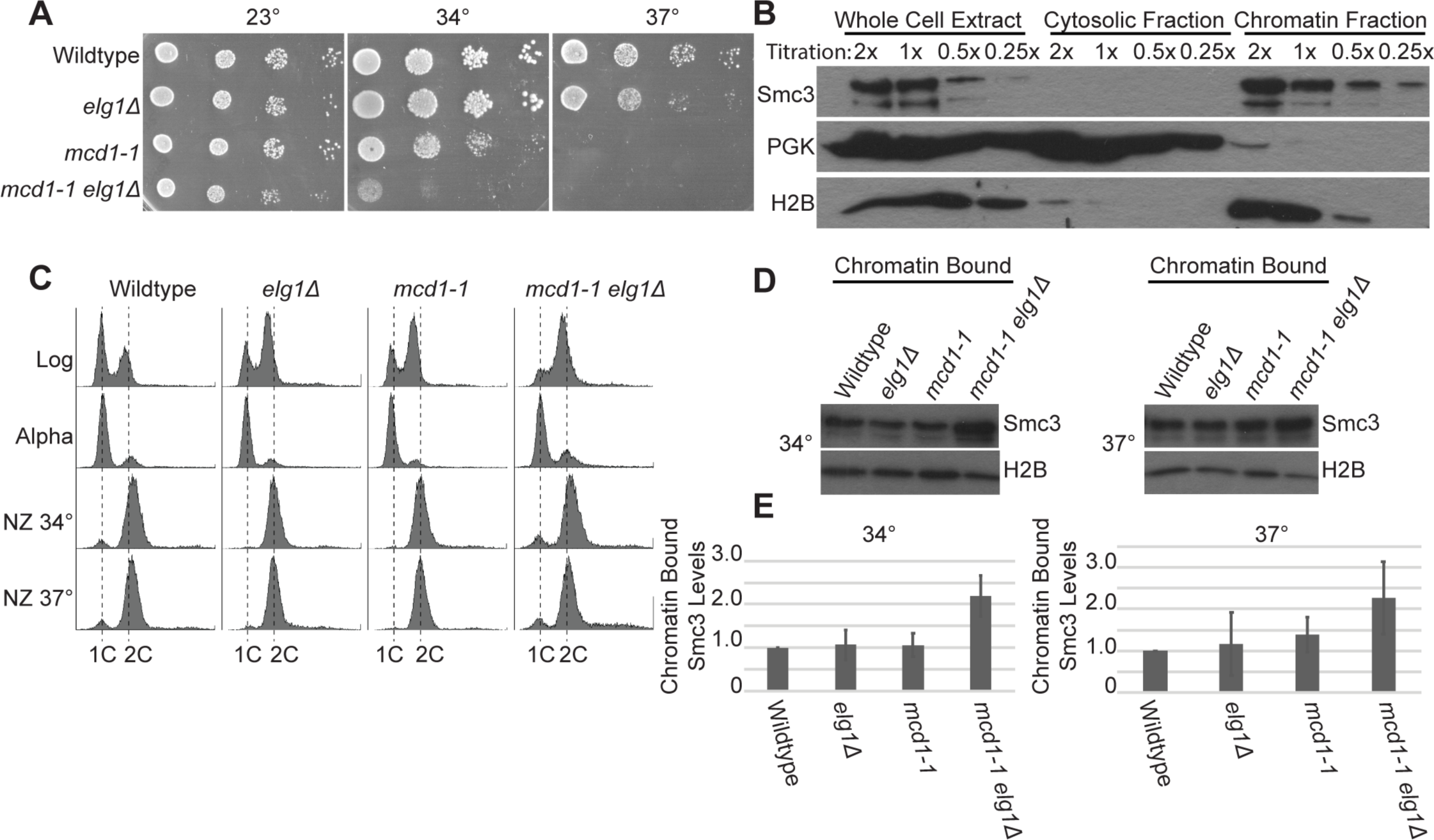
Excess chromatin-bound PCNA (via *elg1Δ*) increases cohesin chromatin levels in *mcd1* mutant cells. [A] 10-fold serial dilution of indicated yeast strains plated on rich medium plates and incubated at 23°C, 34°C, and 37°C for 2 days. [B] Titration and chromatin fractionation of wildtype cells arrested in nocodazole. 1X sample concentration for Smc3, H2B, and PGK indicates samples are within the linear range of detection. [C] DNA content of cells synchronized in G1, and then shifted to 34°C or 37°C and arrested in pre-anaphase. [D] Chromatin bound fraction of cells arrested in pre-anaphase. Smc3 and H2B indicate levels of chromatin-bound proteins in the chromatin fraction. [E] Quantification of chromatin bound Smc3 levels compared to H2B in each noted strain type. Smc3 enrichment on DNA is based on the ratio of Smc3 to Histone 2B levels obtained from 3 independent experiments. Error bars represent standard deviation.

### Elevated levels of chromatin-bound PCNA (via *elg1Δ*) promote Smc3 acetylation in *mcd1* mutant cells

If excess levels of chromatin-bound PCNA (via *elg1Δ*) does not reduce cohesin deposition, PCNA may instead negatively impact acetylation of impaired cohesin complexes to produce growth defects. To test this possibility, wildtype, *elg1Δ* and *mcd1-1* single mutant cells, and *mcd1-1 elg1Δ* double mutant cells, all of which express Smc3-3V5 as the sole source of Smc3 function, were synchronized in G1 and released at 34°C or 37°C into fresh medium supplemented with nocodazole (Figure 2A). Protein samples were then assessed for Smc3 acetylation. Compared to wildtype cells, *elg1Δ* single mutant cells contained similar, if not slightly increased, levels of Smc3 acetylation at both temperatures. In contrast, *mcd1-1* single mutant cells exhibited a dramatic decrease in Smc3 acetylation at both temperatures, despite the retention of Smc3 on DNA (Figure 2B). This provides the first evidence that Mcd1 is critical for Eco1-dependent Smc3 acetylation, but that Mcd1 association with Smc1,3 (even in *mcd1-1* mutant cells, Figure 1) is not sufficient to support acetylation. Importantly, *mcd1-1 elg1Δ* double mutant cells contained increased levels of Smc3 acetylation compared to *mcd1-1* single mutant cells, regardless of a temperature shift to either 34°C or 37°C (Figure 2B, Supplemental Figure 1). This result is consistent with prior evidence that elevated levels of PCNA promote both *eco1* mutant cell viability and Eco1-dependent acetylation of Smc3 [10,15]. In combination, these results reveal that elevated levels of chromatin-bound PCNA, via *elg1Δ*, do not negatively impact either cohesin binding or acetylation.

**Figure 2:**
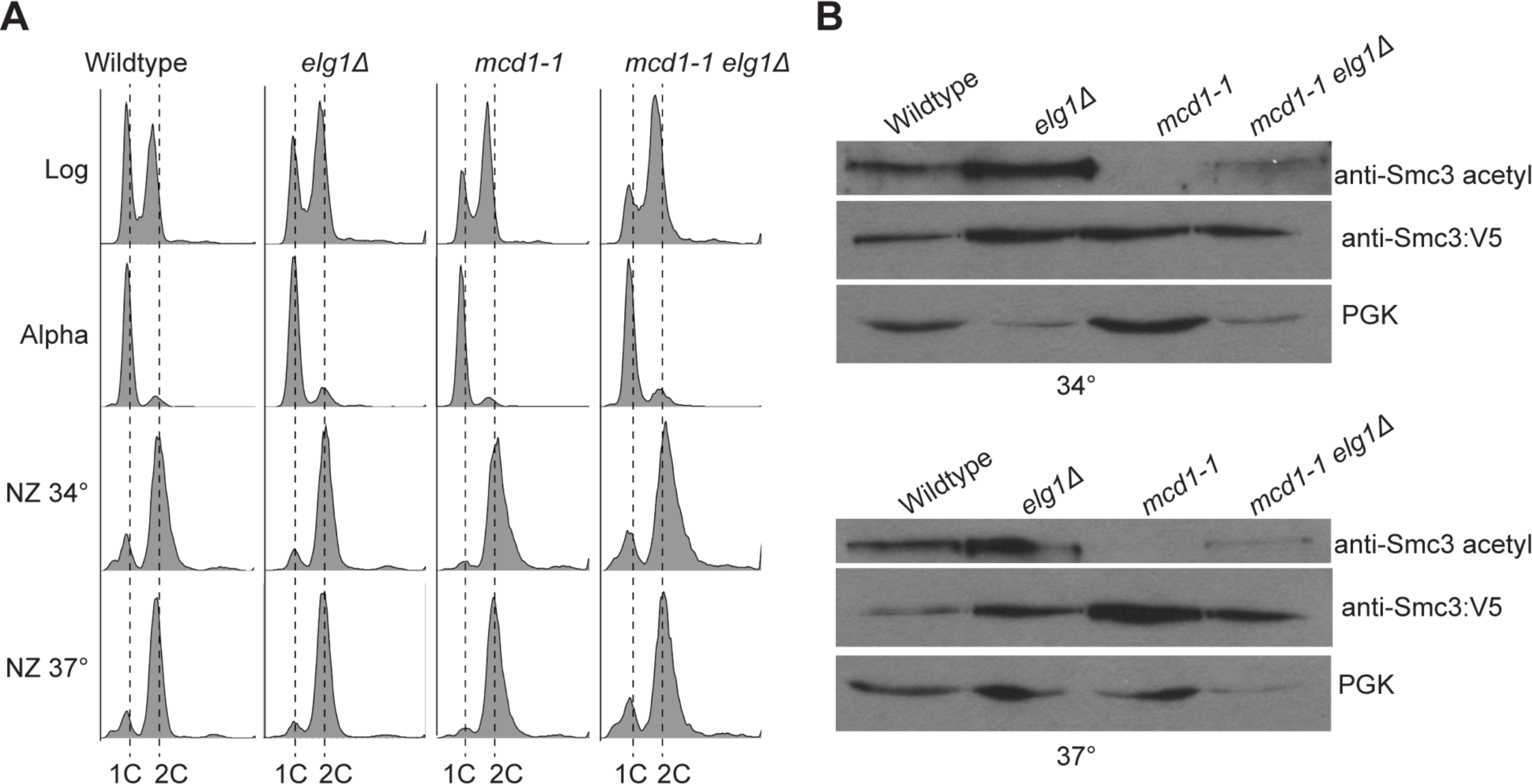
Excess chromatin bound PCNA (via *elg1Δ*) promotes Smc3 acetylation in *mcd1-1* mutant cells. [A] DNA content of log phase cells synchronized in G1, then shifted to 34°C or 37°C and arrested in pre-anaphase. [B] Elevated levels of chromatin-bound PCNA promotes Smc3 acetylation in *mcd1-1* mutant cells. Smc3 was detected by a V5 specific antibody, and acetylated Smc3 was detected by a K112/K113 acetylation antibody. PGK was used as a loading control.

**Supplemental Figure 1: Elevated levels of PCNA promote Eco1-dependent cohesin acetylation.** Second biological iteration of Figure 2. Smc3 was detected by a V5 specific antibody, and acetylated Smc3 was detected by a K112/K113 acetylation antibody. PGK was used as a loading control.

### Elevated levels of chromatin-bound PCNA (via *2μ POL30*) increases *mcd1* mutant cell sensitivity to DNA damage agents

Given the exclusion of models in which elevated levels of chromatin-bound PCNA (via *elg1Δ*) reduce either cohesin binding to DNA or acetylation, we next hypothesized that PCNA-dependent growth defects of cohesin mutants might involve genomic instability. Indeed, it is now well documented that high levels of PCNA, obtained by *GAL*-based overexpression, renders wildtype cells hypersensitive to genotoxic agents and promotes elevated levels of sister chromatid recombination [62], phenotypes similarly exhibited by *elg1Δ* cells [59,62,71–73]. In contrast, studies involving relatively moderate increased levels of PCNA, via 2*µ* plasmids, have thus far failed to produce overt growth defects in wildtype cells [10,15,57,62,74]. We utilized this same 2*μ* plasmid strategy to ascertain whether a modest increase of chromatin-bound PCNA produces increased sensitivity to genotoxic agents in the context of cohesin mutant cells. Serial dilutions of wildtype and *mcd1-1* mutant cells, harboring either *CEN* vector alone, *2μ* vector containing *POL30* (PCNA), were plated on selective media, supplemented with increasing concentrations of MMS, and maintained at the permissive temperature of 23°C. Wildtype cells, with or without the *2μ POL30* plasmid, exhibited similar dose-dependent growth defects, but remained viable even at elevated levels of MMS (Figure 3). *mcd1-1* single mutant cells, that harbored the vector alone, exhibited growth rates nearly identical to those of wildtype cells across the entire dose range of MMS. In contrast, *mcd1-1* mutant cells that harbored the *2μ POL30* plasmid (PCNA^OE^) produced significant growth defects that coincided with increased MMS levels (Figure 3). In combination, this result suggests that even moderate PCNA^OE^ adversely impacts genomic integrity which is synergistic with the effects of MMS and can render *mcd1-1* mutant cells inviable at temperature permissive for *mcd1-1* mutant cells.

**Figure 3:**
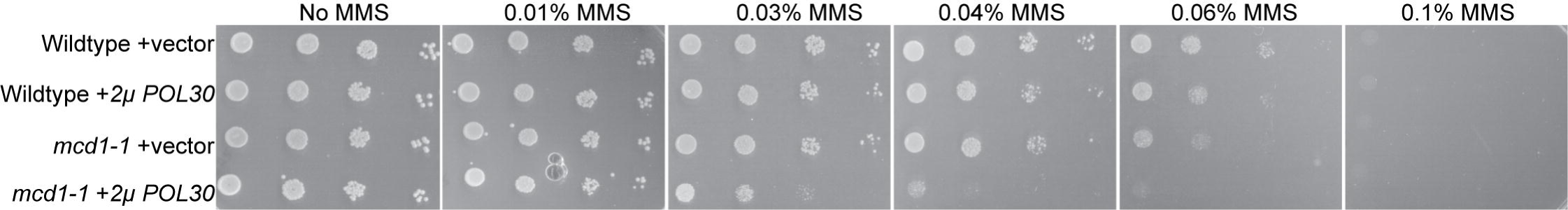
Elevated PCNA levels (via PCNA^OE^) causes DNA damage sensitivity in *mcd1-1* mutant cells. 10-fold serial dilution of indicated yeast strains plated on selective medium plates containing no MMS, 0.01% MMS, 0.03% MMS, 0.04% MMS, 0.06% MMS, or 0.1% MMS, and incubated at 23°C for 3 days.

We next tested whether this moderate PCNA^OE^ is sufficient to sensitize *mcd1-1* mutant cells to replication fork destabilizing agents. Hydroxyurea (HU) inhibits RNR function, resulting in depletion of dNTPs and destabilized DNA replication forks [75]. Serial dilutions of wildtype and *mcd1-1* mutant cells, harboring either *CEN* vector alone or with *2μ POL30*, were exposed to various concentrations of HU and maintained at the permissive temperature of 23°C. The results reveal that PCNA^OE^ produces hypersensitivity to HU even in wildtype cells (Figure 4).

**Figure 4:**
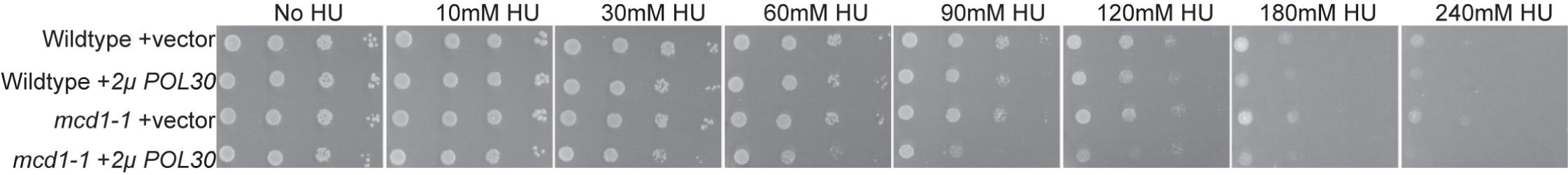
Elevated levels of PCNA sensitizes wildtype and *mcd1-1* mutant cells to HU. 10-fold serial dilution of indicated yeast strains plated on selective medium plates containing no HU, 10mM HU, 30mM HU, 60mM HU, 90mM HU, 120mM HU, 180mM HU, or 240mM HU, and incubated at 23°C for 3 days.

Interestingly, *mcd1-1* mutant cells harboring the vector plasmid alone exhibited growth comparable to that of wildtype cells at all concentrations of HU. *mcd1-1* mutant cells that harbor *2μ POL30* plasmid, however, exhibited significant growth defects even at relatively moderate levels of HU (Figure 4). These results suggest that cohesin plays a critical role in stalled replication fork restart, consistent with prior studies [76–79] and that even moderate increases in chromatin-bound PCNA produces replication stress that is normally remedied in wildtype cells that contain fully functional cohesins.

### Elevated levels of chromatin-bound PCNA (via *2μ POL30*) activate the Mec1/ATR DNA damage response pathway and impact cell cycle progression

The adverse effect of *2μ POL30* on both *mcd1-1* mutant and wildtype cells suggests that even very moderate levels of PCNA is detrimental to genome maintenance. Mec1 (ATR) and Tel1 (ATM) kinases form parallel DNA damage response pathways [80–83]. To determine which, if either, pathway responds to PCNA^OE^, *tel1Δ1* and *mec1-1* single mutant cells were transformed with a *CEN* vector alone or *2µ* vector with *POL30* and grown on plates maintained at a range of temperatures. Surprisingly, neither *tel1Δ1* nor *mec1-1* mutant cells exhibited any adverse growth rates in response to PCNA^OE^, regardless of the temperature (Figure 5A). We next assessed if either *tel1Δ1* or *mec1-1* mutant cells, sensitized by low doses of MMS, might reveal adverse effects of PCNA^OE^. *tel1Δ1* mutant cells that harbor the *2μ POL30* plasmid exhibited growth nearly identical to that of wildtype cells, despite the presence of MMS. Conversely, *mec1-1* mutant cells that harbored the *2μ POL30* plasmid exhibited significant growth defects in the presence of MMS (Figure 5B). These results, and the findings above, suggest that PCNA^OE^ generates genomic instabilities that activate a Mec1/ATR (but not Tel1/ATM) DNA damage repair pathway, and that cohesin is necessary to repair this damage.

**Figure 5:**
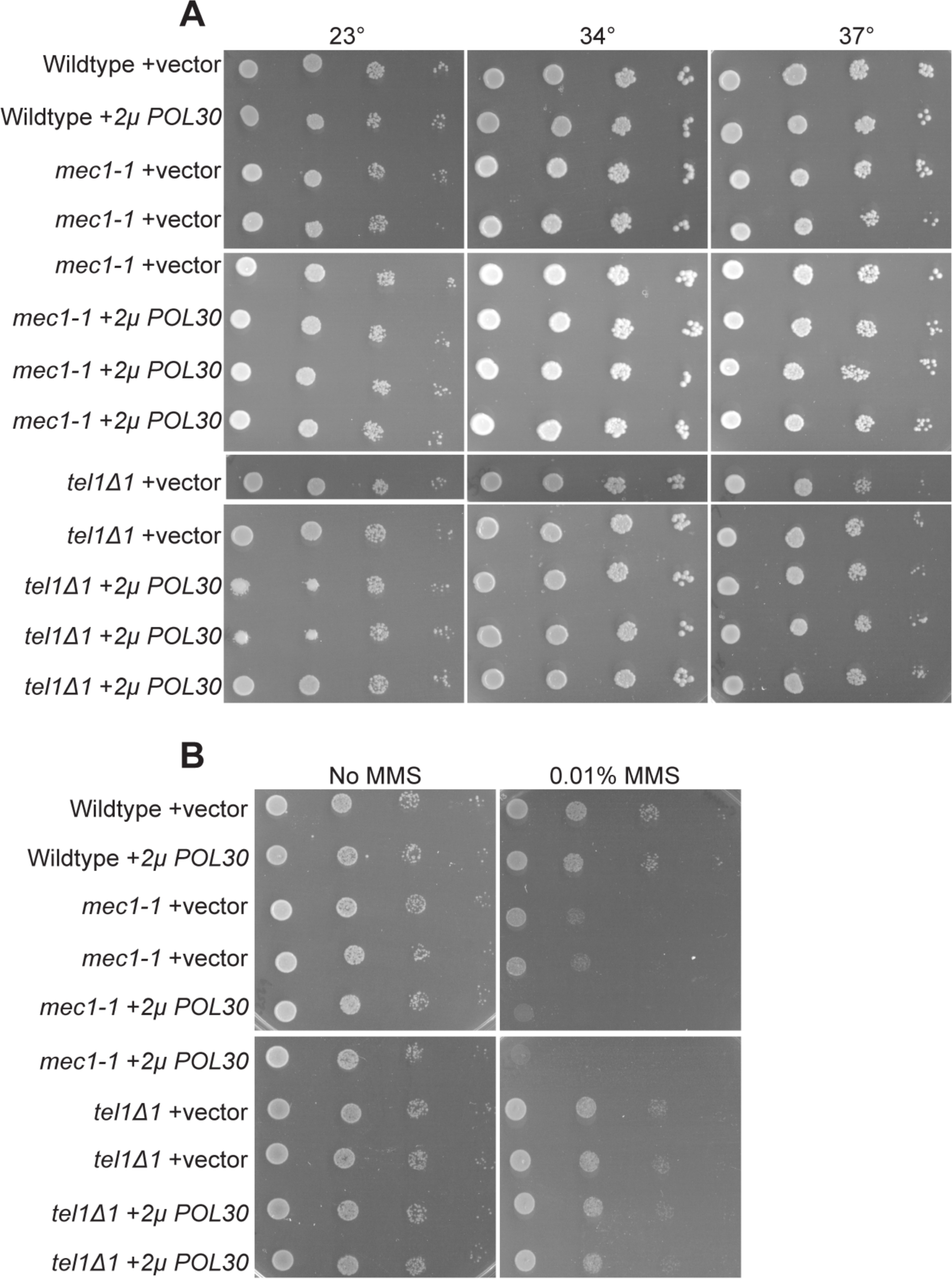
Elevated levels of PCNA sensitizes *mec1-1* mutant cells to MMS. [A] Elevated levels of PCNA does not impact *tel1*Δ*1* and *mec1-1* mutant cells under high temperature stress. 10-fold serial dilution of indicated yeast strains plated on selective medium plates and incubated at 23°C, 34°C, and 37°C for 2 days. [B] Elevated levels of PCNA specifically sensitizes *mec1-1* mutant cells when exposed to DNA damage. 10-fold serial dilution of indicated yeast strains plated on selective medium plates containing no MMS or 0.01% MMS and incubated at 23°C for 2 days.

PCNA^OE^ sensitizes *mcd1-1* mutant cells to MMS and HU and also activates the Mec1/ATR intra-S phase checkpoint, suggesting that the adverse effect imposed by PCNA is through replication stress. Consistent with this hypothesis, mutation of either *ESCO2* (human homolog of yeast *ECO1*) or cohesin subunits result in slowed fork progression in mammalian cells [84–87], although a velocity reduction is absent in analogous yeast mutant strains [2,3,10,76,79,88–90]. It thus became important to test whether elevated levels of chromatin-bound PCNA produces replication stress severe enough to delay S phase progression in *mcd1-1* mutant cells. Log phase wildtype, *elg1Δ, mcd1-1* and *mcd1-1 elg1Δ* double mutant cells were synchronized in G1, washed, and released into fresh medium supplemented with nocodazole and maintained at the either 32° or 37°C (restrictive temperatures for *mcd1-1 elg1Δ* double mutant and *mcd1-1* single mutant cells, respectively). Aliquots were harvested at 15 minute intervals and cell cycle progression monitored by flow cytometry (Supplemental Figure 2). Because cells harboring *ELG1* deletion do not fully synchronize in G1 with alpha factor [31], we analyzed the time point at which at least half of a G1 arrested population progressed to an S-phase state. At 32°C, wildtype and *elg1Δ* mutant cells both reached a half-way point (relatively equal 1N and 2N peaks) between the 30-45 minute time points, while arresting in pre-anaphase in about 75 minutes. *mcd1-1* mutant cells progressed to a mid-way state at the tail end of the time range (45 minutes) required by wildtype and *elg1Δ* mutant cells, and achieved a final preanaphase state in coordination with wildtype and *elg1Δ* mutant cells. At 37°C, which is non-permissive for both *mcd1-1* and *mcd1-1 elg1Δ* mutant cells, *mcd1-1* cells are clearly delayed by almost two time points in progressing to a half-way state (75 minutes), compared to wildtype and *elg1Δ* mutant cells (45-60 minutes) and delayed by a full time point in achieving a preanaphase arrest (90 minutes in *mcd1-1* cells compared to 75 minutes in both wildtype and *elg1Δ* cells. Interestingly, *mcd1-1 elg1Δ* double mutant cells instead both exit G1 and achieve a mid-way state (15 minutes) faster than all other strains at both 32°C and 37°C, but then remain in this mid-way state for an extended period of time of 30 minutes (15-45 minutes) at 37°C, relative to other strains (Supplemental Figure 2). These results suggest that elevated levels of chromatin-bound PCNA, in combination with cohesin mutation, adversely impacts replication fork progression. These findings are consistent with the adverse effects of either HU or MMS in *mcd1-1* PCNA^OE^ cells (Figures 3 and 4, respectively), and suggest that PCNA^OE^-dependent cohesin mutant cell inviability is due to defects in replication stress that resolve into genomic instabilities that accrue over multiple divisions.

**Supplemental Figure 2: Excess chromatin bound PCNA does not cause a S-phase checkpoint activation in wildtype or *mcd1-1* mutant cells.** DNA content of log phase cells synchronized in G1. Temperature was then shifted to either 32°C or 37°C and cells released into media supplemented with nocodazole. Samples were collected every 15 minutes.

### The differential impacts of PCNA^OE^ on various cohesin mutant cells

High levels of PCNA (*GAL*-dependent *POL30* overexpression) results in phenotypes that overlap with those obtained by *elg1Δ*, although *elg1Δ* effects are far more robust and include genomic instabilities that include elevated levels of sister chromatid recombination, chromosomal translocations, fusions, and telomere elongation [62,72,73,91]. It thus became important to ascertain the effect of *elg1Δ* on cohesin mutants to that of a moderate increase (2*µ*-based) in PCNA levels. Our results document that, not only *mcd1-1* cells, but also *smc1-259* and *smc3-42* mutant cells exhibit growth defects in response to *elg1Δ* (Figure 6A), consistent with prior studies [32,34]. We next tested whether wildtype PCNA (via 2*µ* plasmid) would similarly exacerbate cohesin mutant cell growth. The results reveal that PCNA^OE^ severely exacerbates the temperature sensitivity of *mcd1-1* mutant cells (Figure 6B), consistent with a previous study [15]. Surprisingly, PCNA^OE^ exhibited no adverse effect on either *smc1-259* or *smc3-42* mutant cell growth (Figure 6B). We repeated our analysis on different cohesin alleles. The results show that PCNA^OE^ similarly failed to adversely impact the growth of *smc1-2* and *smc3-5* mutant cells. At first blush, these results appear in stark contrast to those reported by Zhang and colleagues for both *smc1-259* and *smc3-42* mutant cell [15]. Our results, however, are predicated on PCNA expressed from the endogenous promoter in which elevated expression is driven solely by the multi-copy 2*µ* plasmid [10]. In contrast, the Zhang study expressed PCNA using the constitutive and high-expressing ADH promotor in the context of the high copy 2*µ* plasmid (pGADT7), suggesting that the two results may differ due to PCNA expression levels. To test this hypothesis, we cloned *POL30* into pGADT7. The resulting PCNA construct, however, produced a dominant-negative phenotype such that all strains tested, including wildtype cells, exhibited severe growth defects (Figure 6C). It is possible that this dominant-negative effect is due to elevated levels of epitope-tagged PCNA via pGADT7, tagging which is absent in our 2*µ* strategy. Prior studies, for instance, document that endogenous expression of either N- and C-terminus tagged PCNA can result in wildtype cell increased MMS sensitivity [61,74]. Regardless, our results reveal that *mcd1-1*, in contrast to numerous other cohesin subunit alleles (which exhibit similar conditional growth defects - Figure 6), are hypersensitive to even a moderate increase in wildtype PCNA.

**Figure 6:**
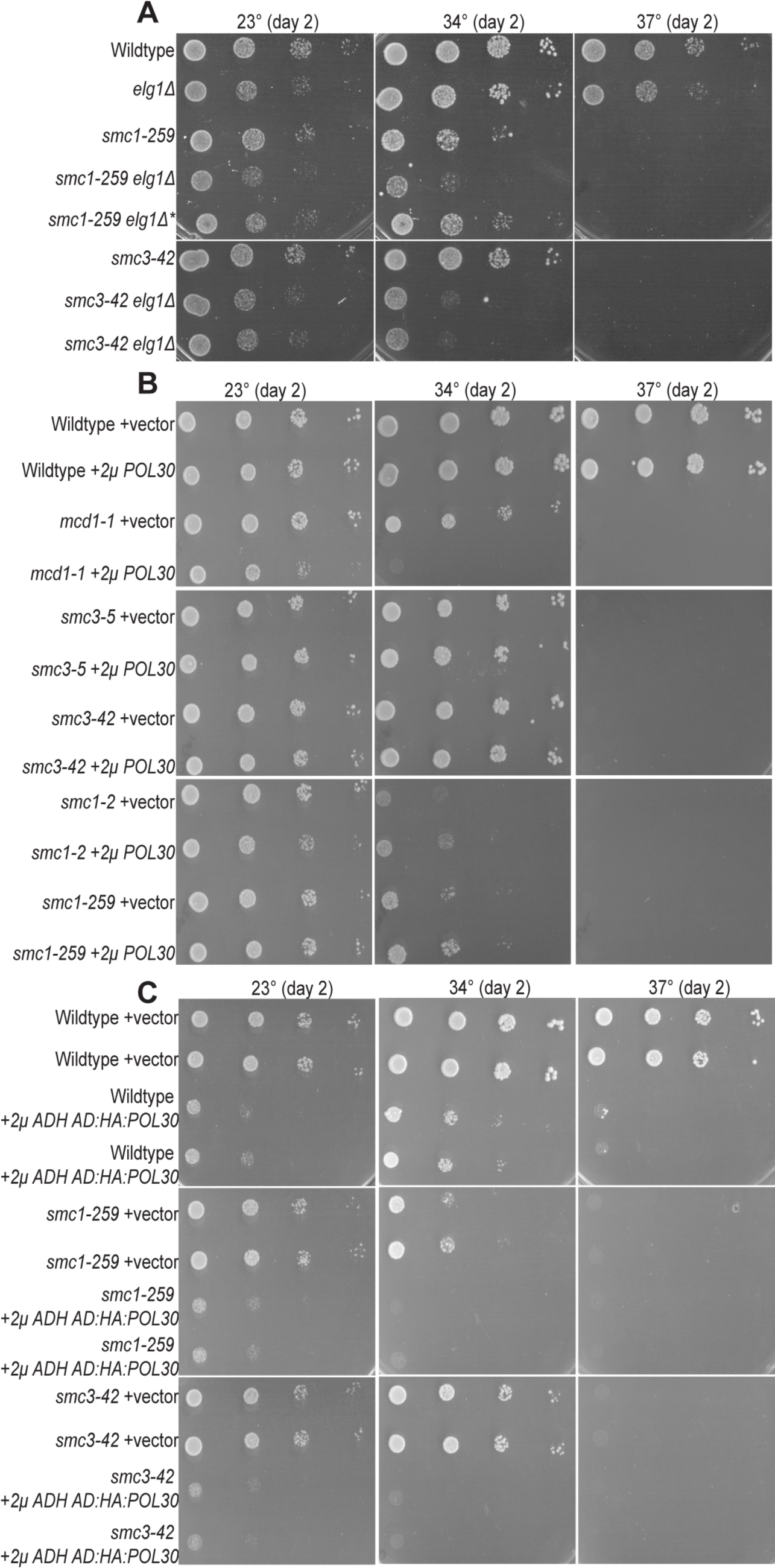
Elevated levels of chromatin-bound PCNA differentially impacts cohesin alleles. [A] *ELG1* deletion exacerbates *smc1-259* and *smc3-42* mutant cells. 10-fold serial dilution of indicated yeast strains plated on rich medium plates and incubated at 23°C, 34°C, and 37°C for 2 days. * indicates a mutated *smc1-259 elg1Δ* strain that displays a resistant to temperature sensitivity [B] PCNA^OE^ from a 2μ plasmid specifically impacts *mcd1-1* mutant cells. 10-fold serial dilution of indicated yeast strains plated on selective medium plates and incubated at 23°C, 34°C, and 37°C for 2 days. [C] Overexpressed PCNA harboring a GAL4 AD and HA N-terminal tag exacerbates cell growth in wildtype cells, *smc1-259*, and *smc3-42* mutant cells. Images of *POL30* in pGADT7 in WT, *smc3-42*, and *smc1-259* mutant cells. 10-fold serial dilution of indicated yeast strains plated on selective medium plates and incubated at 23°C, 34°C, and 37°C for 2 days.

## DISCUSSION

Since the first evidence that cohesion establishment is intimately coupled to DNA replication [10], a wealth of studies document the dependency of Eco1 recruitment by DNA replication fork factors and, furthermore, that replication fork factors regulate sister chromatid tethering reactions [14,15,22–24,28,31,32,34,35,92–96]. Of particular interest are RFC complexes that regulate the loading (RFC1^RFC^, Ctf18^RFC^) and unloading (Elg1^RFC^) of PCNA onto DNA [60,97–99]. Either PCNA overexpression or *ELG1* deletion, both of which result in prolonged and elevated retention of PCNA onto DNA, rescue *eco1* mutant cell growth [10,15,31,32]. Intriguingly, however, elevated levels of chromatin-bound PCNA instead exacerbates the growth defects of the cohesin subunits [15,32], a surprising phenotype given that the cohesin subunit Smc3 is the target of Eco1 acetylation [12,13,100]. *ELG1* deletion, however, is mutagenic and thus may have adverse impacts beyond PCNA unloading while moderate increases in PCNA appear fully tolerated by wildtype cells [57,59,62,72,74,101]. Here, we confirm that *ELG1* deletion exacerbates the growth defects present in *mcd1* mutant cells and extend those findings to show that simply moderate overexpression of wildtype PCNA similarly exerts an adverse growth effect on *mcd1* mutant cell growth. The link between PCNA and cohesin function is complex, however, in that similar expression of PCNA had no impact on a number of other cohesin subunit alleles (*smc3-5, smc3-42, smc1-295* and *smc1-2*). These findings raise the possibility that cohesin loading onto sister chromatids, that occurs immediately behind DNA polymerase and PCNA, may occur in an orientation in which Mcd1 is most proximal to, and thus most impacted by, DNA replication fork components. Given that deletion of the PCNA-unloader component Elg1 exacerbates *mcd1* mutant cell growth, a simple prediction is that deletion of the PCNA RFC loader component Ctf18 should suppress cohesin mutant cell growth defects. Instead, *CTF18* deletion is lethal, or produces significant growth defects, in *smc1-259* mutant cells [102] and also in *eco1* mutant cells [10]. Clearly, additional studies are required to elucidate the multiple mechanisms through which DNA replication fork components, especially involving PCNA, impact cohesin functions.

A second major finding from this study is the exclusion of likely mechanisms through which elevated levels of chromatin-bound wildtype PCNA adversely impacts cohesin functions. Elevated levels of chromatin-bound PCNA that persist well after S phase could antagonize any number of post-replicative activities such as histone deposition, chromatin remodeling, or Okazaki maturation [103,104]. Given the adverse impact of elevated levels of chromatin-bound PCNA on *mcd1-1* mutant cells, PCNA might conceivably reduce Scc2,4 recruitment and/or subsequent deposition of cohesins onto DNA [8,22,65,105,106]. An alternate model is that elevated levels of PCNA sequester Eco1 into soluble PCNA pools, precluding sufficient cohesin acetylation. We provide evidence that negate both of these models. Our finding that elevated chromatin-binding of PCNA actually improves Smc3 acetylation is consistent with prior findings [10,15]. More importantly, the finding that increased PCNA promotes cohesin binding onto DNA suggests that factors that promote cohesin deposition (Scc2,4, Chl1, or RSC), or promote cohesin dissociation (Rad61), are themselves sensitive to PCNA levels or modifications [8,22,23,58,96]. In support of this model are reports that PCNA physically interacts with Chl1, which promotes the recruitment to DNA of both Scc2,4 and cohesin [14,22,65,106]. Intriguingly, increased Smc3 acetylation, induced by *elg1Δ*, fails to rescue *mcd1-1* mutant cell growth defects, suggesting that PCNA impacts cohesin functions beyond sister chromatid tethering pathways - as described below.

A third major finding from the current study involves identifying the mechanism through which PCNA renders *mcd1-1* mutant cells inviable. It is well known that *elg1Δ* results in long-lived and elevated levels of chromatin-bound PCNA which is highly mutagenic to wildtype cells [72,73,91]. High levels of PCNA, achieved by *GAL*-overexpression or coupling *ADH*-overexpression to 2*µ*-based high copy number, similarly produce genomic instabilities [15,62]. A much more moderate expression system (2*µ* high copy) has thus far shown no adverse effect in wildtype cells [10,57]. Our findings suggest that even this moderate form of increased expression of PCNA produces genomic instability, an effect revealed in wildtype cells only by synergistic challenging with MMS or HU. PCNA^OE^ is sufficient to produce growth defects in *mcd1-1* cohesin mutants, and this effect is greatly exacerbated by challenging mutant cells with MMS or HU. This intersection between PCNA and cohesin is consistent with prior studies that document that 1) highly elevated levels of PCNA result in genomic instability [62], 2) cohesin are both recruited to stalled replication forks and promote fork restart [76–78,87], and 3) DNA replication fork protection complexes and cohesin pathways are intimately linked [23,62,76,79,107]. During the final stages of this study, independent analyses revealed that elevated levels of chromatin-bound PCNA results in hyper-recruitment of mismatch repair factors [108]. Cells that exhibit elevated levels of mismatch repair intermediates normally activate cohesin recruitment pathways to promote efficient DNA repair through homologous recombination [109,110]. While the moderate increase of PCNA used here does not appear to generate long-lived genomic instabilities sufficient to require mismatch repair factor Msh3 (Supplemental Table 4). Cells utilize Msh2-Msh3-dependent mismatch repair (MMR) to correct mismatches (including short insertions and deletions) that accumulate during DNA replication [108,111–113], which increase upon high levels of PCNA expression [108]. However, numerous *mcd1-1 msh3Δ elg1Δ* triple mutant cells were obtained from crossing *msh3Δ* single mutant cells to *mcd1-1 elg1Δ* double mutant cells (Supplemental Table 4). Despite this, our results clearly document that the adverse effect of PCNA^OE^ does activate a Mec1(ATR)-dependent response, versus a Tel1(ATM)-dependent response. The integration of Mec1 sensor in PCNA and cohesin genome instability contrasts those in which double deletion of *MEC1* and *TEL1* were required to reduce Scc1 recruitment to forks under conditions of replication stress [76]. In combination, these findings highlight the complex nature through which Mec1 (and possibly Tel1) respond to replication stressors (such as PCNA^OE^), which are revealed only under conditions of cohesin mutation and/or genotoxic agents.

A confluence of findings document that 1) cohesins are deposited during S phase onto nascent sister chromatids [8,10,64,114–119], 2) cohesins are critical for replication fork restart and repair [76–78,86], 3) Eco1-dependent cohesion establishment is intimately coupled to PCNA and the DNA replication fork [10,15,16,24–26,120–123], and 4) PCNA^OE^ and *elg1Δ* both adversely impact cohesin mutant cell growth. Clearly, PCNA^OE^ (at even very moderate levels) and *elg1Δ* adversely impact cohesin mutant cell growth, an effect exacerbated by genotoxic agents. We thus envision two scenarios for these effects, both of which are predicated on PCNA^OE^-dependent replication stress (Figure 7). One scenario involves competition for Eco1 activity. For instance, Eco1 modifies PCNA during DNA damage [124]. Elevated levels of chromatin-bound PCNA, along with Smc3 acetylation, may outcompete the ability of Eco1 to modify the appropriate pool of PCNA when the fork is under stress due to PCNA^OE^. Competition for Eco1 activity may further involve Mcd1. While Smc3 is acetylated during DNA replication to promote sister chromatid cohesion, Eco1 instead modifies Mcd1 in response to DNA damage [125,126]. In the future, it will be important to test the extent to which Smc3 and retained PCNA may outcompete Mcd1 acetylation during replication stress, a model consistent with our finding that *mcd1-1* mutant cells are highly sensitive to PCNA mis-regulation, relevant to other cohesin mutant strains. An alternative scenario posits that the increase of chromatin-bound cohesin in response to PCNA^OE^, reduces replication, resulting in increased genomic instability and cell death [56,77,79,87]. These findings may provide new insights through which cancer cells that harbor cohesin gene mutations [127–132] can be targeted.

**Figure 7:**
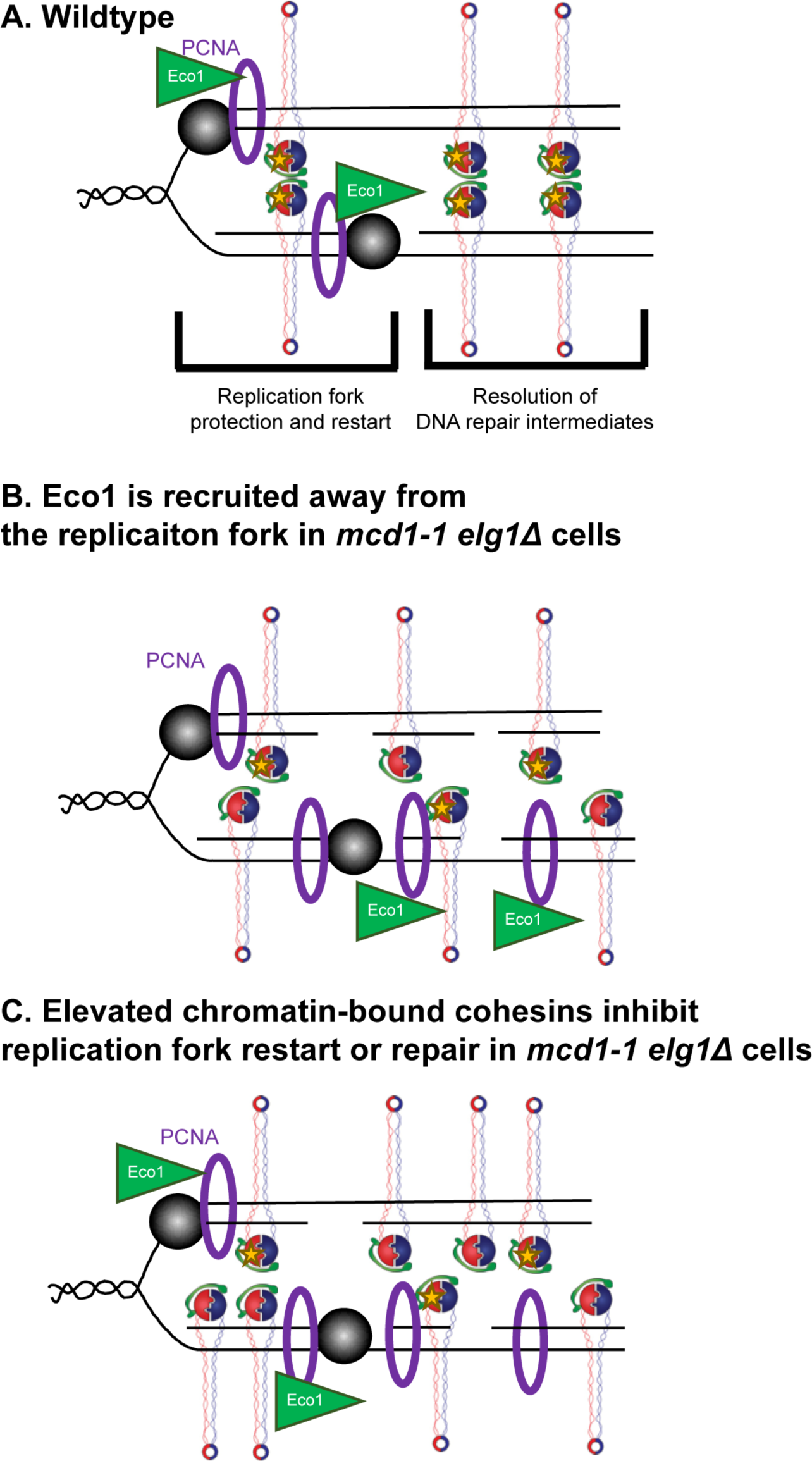
Cohesins are necessary to repair and restart stalled replication forks caused by elevated levels of PCNA. [A] Stalled replication forks result in DNA damage and the emergence of DNA repair intermediates. Through Mec1-dependent signaling, wildtype cells repair and restart the replication fork, and resolve DNA repair intermediates and damage [B]. In *mcd1-1 elg1Δ* double mutant cells, retained chromatin-bound PCNA may recruit Eco1 away from the replication fork to target Smc3, therefore reducing PCNA modification essential in DNA damage repair. [C] An alternate model that the elevated levels of chromatin-bound cohesin hinder replication fork restart and repair in *mcd1-1 elg1Δ* double mutant cells, leading to cell death.

**Supplemental Table 4: *msh3Δ* does not result in cell spore lethality in *mcd1-1 elg1Δ* double mutant cells.** Dissection of *mcd1-1 elg1Δ Smc3:3V5* mated with *msh3Δ*. *mcd1-1 elg1Δ msh3Δ* triple mutant strains are obtained at the expected frequency.

## ACKNOWLEDGEMENTS

We thank the R.V.S. laboratory members (Michael Mfarej, Annie Sanchez, Nicole Kirven, and Shaya Ameri) for helpful comments during the progression of this study. We also thank Dr. Vincent Guacci, Dr. Douglas Koshland, Dr. Gregory Lang, and Dr. Katsuhiko Shirahige for generously sharing yeast strains and reagents. Any opinions, findings, and conclusions or recommendations expressed in this study are those of the authors and does not necessarily reflect the views of the National Institutes of Health. No competing interests are declared.

