## Supplementary figures and images for "PCNA Antagonizes Cohesin-dependent Roles in Genomic Stability"

### Supplemental Figure 1.tif

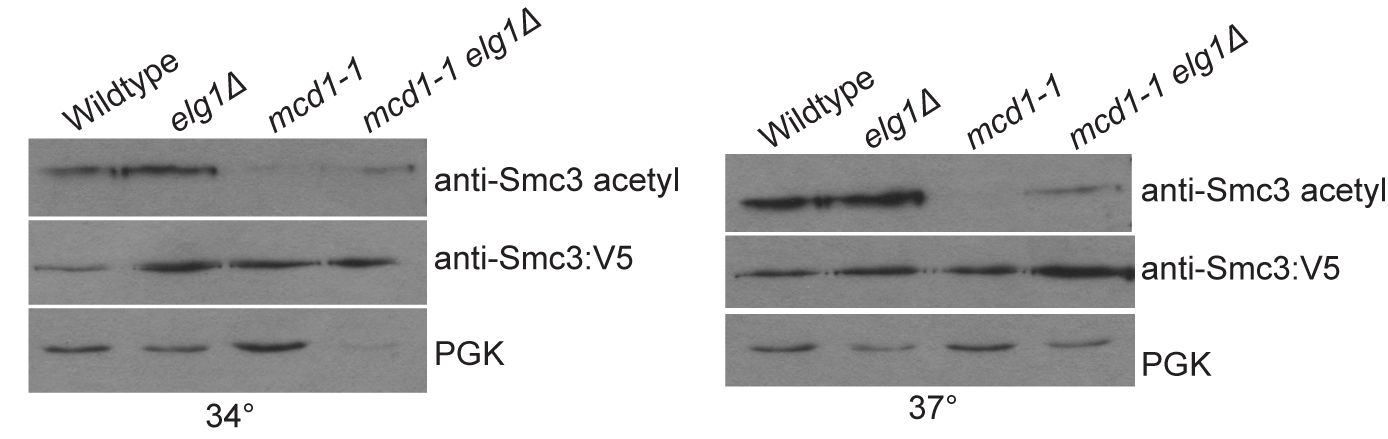

### Supplemental Figure 2.tif

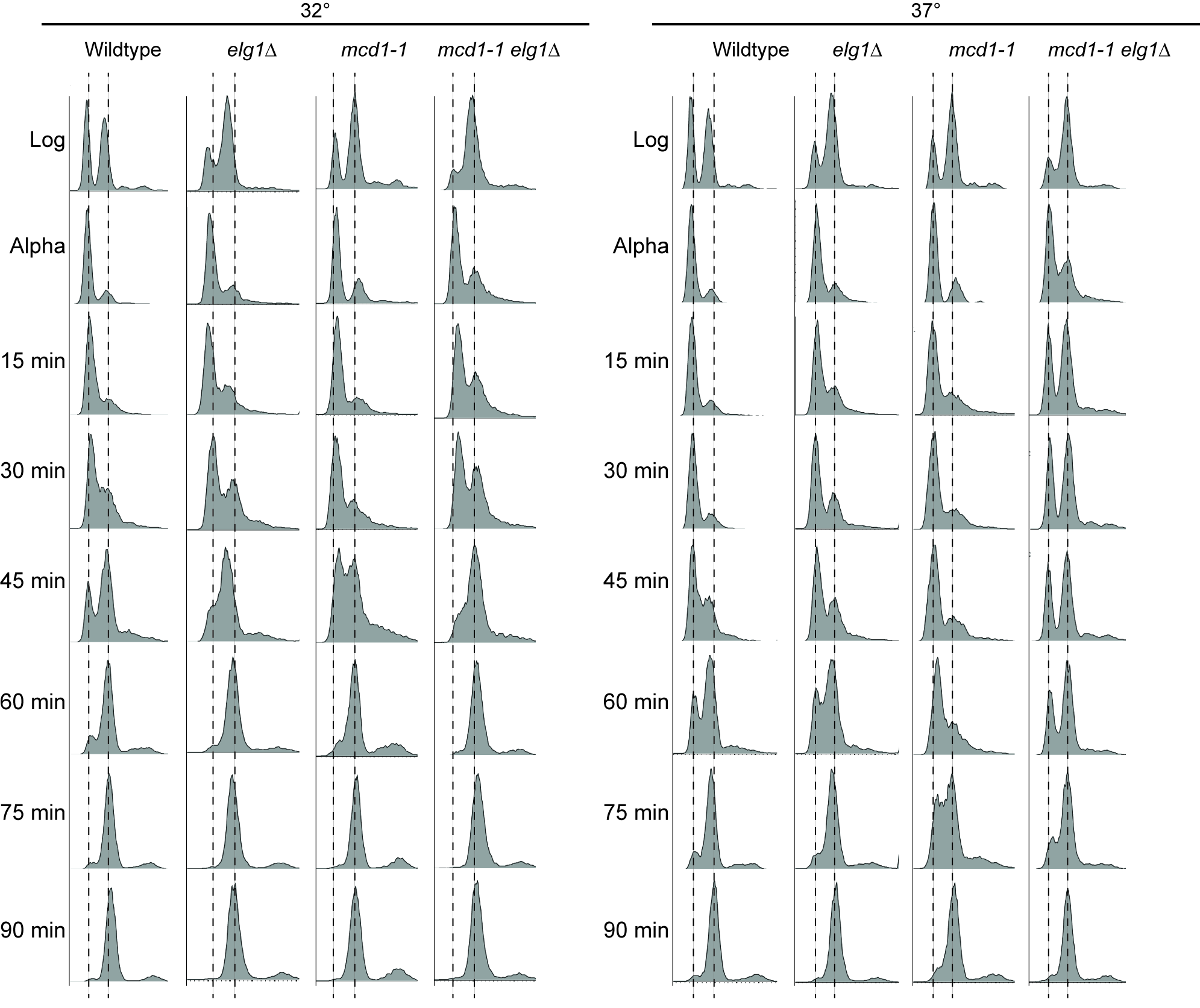
