## Supplemental Tables 1-4 for "PCNA Antagonizes Cohesin-dependent Roles in Genomic Stability"

| Supplemental Table 1: Reagents used in this study | | | | |
| --- | --- | --- | --- | --- |
| Reagent type | **Designation** | **Source or reference** | **Identifiers** | **Additional Information** |
| Chemical compound | Alpha Factor | Zymo | Y1001 |  |
| Chemical compound | Nocodozole | Sigma | M1404 |  |
| Chemical compound | Methyl methanesulfonate | ACROS | 66-27-3 | noted as MMS in text |
| Chemical compound | Hydroxyurea | Sigma | H8627 | noted as HU in text |
| Chemical compound | Protease inhibitor cocktail | Sigma | P8215 |  |
| Chemical compound | ECL Prime | GE Healthcare | RPN2232 |  |
| Chemical compound | Glass beads | BioSpec | 11079105 |  |
| Chemical compound | IGEPAL-630 | Sigma | I-3021 |  |
| Antibody | Mouse anti-PGK | Novex | 459250 | WB: 1:20,000 |
| Antibody | Rabbit anti-H2B | Abcam | ab188291 | WB: 1:80,000 |
| Antibody | Mouse anti-V5 | Invitorgen | R960-25 | WB: 1:40,000 |
| Antibody | Mouse anti-SMC3 K112/K113 acetylation | Dr. Katsuhiko Shirahige |  | WB: 1:1,000 |
| Antibody | Goat anti-rabbit | Bio-Rad | 170-6515 | WB: 1:40,000 |
| Antibody | Goat anti-mouse | Bio-Rad | 170-6516 | WB: 1:10,000 (anti-acetyl) or 1:40,000 (V5 and PGK) |
| Genetic Reagent | *Saccharomyces cerevisiae* | this paper | Yeast strains | Supplementary table 2 |
| Genetic Reagent | *Escherichia coli* plasmids | this paper | DNA Plasmids | Supplementary table 3 |

Supplemental Table 2: Strains used in this study

Strain	Genotype	Reference

DLY285	*MATa mec1-1:HIS ura3 leu2 trp1 his3*	*Paulovich, 1995*

EU3430-9A *MATa SMC3-3V5-HIS3MX leu2-3, 112 his3-11, 15 lys2-801 trp1-1 bar1 GAL+*	*Unal, 2008*

K699	*MATa ade 2-1 his 3-11,15 leu 2-3,112 trp 1-1 ura3 can1-100 GAL psi+*	*Schwob, 1994*

K5824	*MATa ade 2-1 his 3-11,15 leu 2-3,112 trp 1-1 ura3 can1-100 smc3-42*	*Stead, 2003*

K6013	*MATa ade 2-1 his 3-11,15 leu 2-3,112 trp 1-1 ura3 can1-100 GAL psi+ smc1-259*	*Stead, 2003*

YBS2021	*MATa ade 2-1 his 3-11,15 leu 2-3,112 trp 1-1 ura3-1 mcd1-1 Net1:GFP:KAN*	This study

YDM884	*MATa tel1∆1::HIS3 ura3-52 his3∆200 ade2-101 leu2∆1 lys2-801 trp1∆1	Morrow, 1995*

YMM433	*MATa ade 2-1 his 3-11,15 leu 2-3,112 trp 1-1 ura3-1 can1-100 smc1-2*	This study

YMM435	*MATa ade 2-1 his 3-11,15 leu 2-3,112 trp 1-1 ura3-1 can1-100 smc3-5*	This study

YCZ044	*MATα ade 2-1 his 3-11,15 leu 2-3,112 trp 1-1 ura3-1*	This study

YCZ143	*MATα ade 2-1 his 3-11,15 leu 2-3,112 trp 1-1 ura3-1 mcd1-1*	This study

YCZ144	*MATa ade 2-1 his 3-11,15 leu 2-3,112 trp 1-1 ura3-1 mcd1-1*	This study

YCZ147	*MATa ade 2-1 his 3-11,15 leu 2-3,112 trp 1-1 ura3-1*	This study

YCZ225	*MATa ade 2-1 his 3-11,15 leu 2-3,112 trp 1-1 ura3-1 mcd1-1 2µ POL30:URA*	This study

YCZ249	*MATa ade 2-1 his 3-11,15 leu 2-3,112 trp 1-1 ura3-1 KAN:ECO1, elg1∆::TRP*	This study

YCZ262	*MATα ade 2-1 his 3-11,15 leu 2-3,112 trp 1-1 ura3-1 SMC3:V5:HIS*	This study

YCZ280	*MATα ade 2-1 his 3-11,15 leu 2-3,112 trp 1-1 ura3-1 elg1∆::TRP SMC3:V5:HIS*	This study

YCZ407	*MATa ade 2-1 his 3-11,15 leu 2-3,112 trp 1-1 ura3-1 SMC3:V5:HIS*	This study

YCZ408	*MATa ade 2-1 his 3-11,15 leu 2-3,112 trp 1-1 ura3-1 mcd1-1 elg1∆::TRP SMC3:V5:HIS*	This study

YCZ421	*MATα ade 2-1 his 3-11,15 leu 2-3,112 trp 1-1 ura3-1 mcd1-1 elg1∆::TRP SMC3:V5:HIS*	This study

YCZ425	*MATa ade 2-1 his 3-11,15 leu 2-3,112 trp 1-1 ura3-1 mcd1-1 SMC3:V5:HIS*	This study

YCZ428	*MATa ade 2-1 his 3-11,15 leu 2-3,112 trp 1-1 ura3-1 elg1∆::TRP SMC3:V5:HIS*	This study

YCZ465	*MATa ade 2-1 his 3-11,15 leu 2-3,112 trp 1-1 ura3-1 2µ POL30:URA*	This study

YCZ474	*MATa ade 2-1 his 3-11,15 leu 2-3,112 trp 1-1 ura3-1 mcd1-1 2µ vector:URA*	This study

YCZ477	*MATa ade 2-1 his 3-11,15 leu 2-3,112 trp 1-1 ura3-1 CEN vector:URA*	This study

YCZ530	*MATa ade 2-1 his 3-11,15 leu 2-3,112 trp 1-1 ura3-1 can1-100 smc1-2 CEN vector:URA*	This study

YCZ532	*MATa ade 2-1 his 3-11,15 leu 2-3,112 trp 1-1 ura3-1 can1-100 smc1-2 2µ POL30:URA*	This study

YCZ534	*MATa ade 2-1 his 3-11,15 leu 2-3,112 trp 1-1 ura3-1 can1-100 smc3-5 CEN vector:URA*	This study

YCZ536	*MATa ade 2-1 his 3-11,15 leu 2-3,112 trp 1-1 ura3-1 can1-100 smc3-5 2µ POL30:URA*	This study

YCZ559	*MATa ade 2-1 his 3-11,15 leu 2-3,112 trp 1-1 ura3 can1-100 smc3-42 CEN vector:URA*	This study

YCZ561	*MATa ade 2-1 his 3-11,15 leu 2-3,112 trp 1-1 ura3 can1-100 smc3-42 2µ POL30:URA*	This study

YCZ563	*MATa ade 2-1 his 3-11,15 leu 2-3,112 trp 1-1 ura3 can1-100 GAL psi+ smc1-259 CEN vector:URA*	This study

YCZ565	*MATa ade 2-1 his 3-11,15 leu 2-3,112 trp 1-1 ura3 can1-100 GAL psi+ smc1-259 2µ POL30:URA*	This study

YCZ567	*MATa mec1-1:HIS ura3 leu2 trp1 his3 CEN vector:URA*	This study

YCZ568	*MATa mec1-1:HIS ura3 leu2 trp1 his3 CEN vector:URA*	This study

YCZ575	*MATa mec1-1:HIS ura3 leu2 trp1 his3 CEN vector:URA*	This study

YCZ569	*MATa mec1-1:HIS ura3 leu2 trp1 his3 2µ POL30:URA*	This study

YCZ570	*MATa mec1-1:HIS ura3 leu2 trp1 his3 2µ POL30:URA*	This study

YCZ576	*MATa mec1-1:HIS ura3 leu2 trp1 his3 2µ POL30:URA*	This study

YCZ571	*MATa tel1∆1::HIS3 ura3-52 his3∆200 ade2-101 leu2∆1 lys2-801 trp1∆1 CEN vector:URA*	This study

YCZ572	*MATa tel1∆1::HIS3 ura3-52 his3∆200 ade2-101 leu2∆1 lys2-801 trp1∆1 CEN vector:URA*	This study

YCZ577	*MATa tel1∆1::HIS3 ura3-52 his3∆200 ade2-101 leu2∆1 lys2-801 trp1∆1 CEN vector:URA*	This study

YCZ573	*MATa tel1∆1::HIS3 ura3-52 his3∆200 ade2-101 leu2∆1 lys2-801 trp1∆1 2µ POL30:URA*	This study

YCZ574	*MATa tel1∆1::HIS3 ura3-52 his3∆200 ade2-101 leu2∆1 lys2-801 trp1∆1 2µ POL30:URA*	This study

YCZ578	*MATa tel1∆1::HIS3 ura3-52 his3∆200 ade2-101 leu2∆1 lys2-801 trp1∆1 2µ POL30:URA*	This study

YCZ662	*MATa ade 2-1 his 3-11,15 leu 2-3,112 trp 1-1 ura3-1 2µ ADH:GAL4AD:HA Vector:LEU*	This study

YCZ663	*MATa ade 2-1 his 3-11,15 leu 2-3,112 trp 1-1 ura3-1 2µ ADH:GAL4AD:HA Vector:LEU*	This study

YCZ664	*MATa ade 2-1 his 3-11,15 leu 2-3,112 trp 1-1 ura3-1 2µ ADH:GAL4AD:HA:POL30:LEU*	This study

YCZ665	*MATa ade 2-1 his 3-11,15 leu 2-3,112 trp 1-1 ura3-1 2µ ADH:GAL4AD:HA:POL30:LEU*	This study

YCZ666	*MATa ade 2-1 his 3-11,15 leu 2-3,112 trp 1-1 ura3-1 smc1-259 2µ ADH:GAL4AD:HA Vector:LEU*	This study

YCZ667	*MATa ade 2-1 his 3-11,15 leu 2-3,112 trp 1-1 ura3-1 smc1-259 2µ ADH:GAL4AD:HA Vector:LEU*	This study

YCZ668	*MATa ade 2-1 his 3-11,15 leu 2-3,112 trp 1-1 ura3-1 smc1-259 2µ ADH:GAL4AD:HA:POL30:LEU*	This study

YCZ669	*MATa ade 2-1 his 3-11,15 leu 2-3,112 trp 1-1 ura3-1 smc3-42 2µ ADH:GAL4AD:HA:POL30:LEU*	This study

YCZ670	*MATa ade 2-1 his 3-11,15 leu 2-3,112 trp 1-1 ura3-1 smc3-42 2µ ADH:GAL4AD:HA Vector:LEU*	This study

YCZ671	*MATa ade 2-1 his 3-11,15 leu 2-3,112 trp 1-1 ura3-1 smc3-42 2µ ADH:GAL4AD:HA Vector:LEU*	This study

YCZ672	*MATa ade 2-1 his 3-11,15 leu 2-3,112 trp 1-1 ura3-1 smc3-42 2µ ADH:GAL4AD:HA:POL30:LEU*	This study

YCZ673	*MATa ade 2-1 his 3-11,15 leu 2-3,112 trp 1-1 ura3-1 smc3-42 2µ ADH:GAL4AD:HA:POL30:LEU*	This study

YCZ702	*MATa/α ade 2-1 his 3-11,15 leu 2-3,112 trp 1-1 ura3-1 smc1-259 elg1∆::KAN*	This study

YCZ706	*MATa ade 2-1 his 3-11,15 leu 2-3,112 trp 1-1 ura3-1 smc3-42 elg1∆::KAN*	This study

YCZ707	*MATa ade 2-1 his 3-11,15 leu 2-3,112 trp 1-1 ura3-1 smc3-42 elg1∆::KAN*	This study

YCZ728	*MATa ade 2-1 his 3-11,15 leu 2-3,112 trp 1-1 ura3-1 smc1-259 elg1∆::KAN*	This study

YCZ730	*MATα ade 2-1 his 3-11,15 leu 2-3,112 trp 1-1 ura3-1 smc1-259 elg1∆::KAN*	This study

YCZ778	*MATa his3-1 leu2-0 lys2-0 ura3-0 msh3∆::KAN	Baudin, 1993* *and Wach, 1994*

YCZ799	*MATa/α ade 2-1 his 3-11,15 leu 2-3,112 trp 1-1 ura3-1 his3-1 leu2-0 lys2-0 ura3-0 msh3∆::KAN elg1∆::TRP mcd1-1*	This study

All strains are in the W303 background except for DLY285 (A364a), EU3430-9A (A364a), YDM884 (A364a), and YCZ778 (S288C) and YCZ799 (W303/S288C diploid)

Supplemental Table 3: Plasmids used in this study

Plasmid	Genotype	Reference

pBS99	*2u POL30:URA*	*Skibbens et al., 1999*

pRS316	CEN Vector *CEN:URA*	*Sikorski,. and Hieter., 1989*

pGADT7	*2µ* Vector *ADH:HA:LEU*	*TakaraBio #630442*

pCZ058	*2µ* *ADH:HA:POL30:LEU*	*This study*

| Supplemental Table 4: Yeast tetrad dissections of *mcd1-1 elg1Δ Smc3:3V5* x *msh3Δ* | | |
| --- | --- | --- |
| Genotype | **Expected** | **Observed** |
| Wildtype | 5 | 0 |
| *mcd1-1* | 5 | 5 |
| *elg1Δ* | 5 | 3 |
| *msh3Δ* | 5 | 0 |
| Smc3:3V5 | 5 | 1 |
| *mcd1-1 elg1Δ* | 5 | 4 |
| *mcd1-1* Smc3:3V5 | 5 | 2 |
| *mcd1-1 msh3Δ* | 5 | 0 |
| *elg1Δ* Smc3:3V5 | 5 | 3 |
| *elg1Δ msh3Δ* | 5 | 3 |
| *msh3Δ* Smc3:3V5 | 5 | 0 |
| *mcd1-1 elg1Δ* Smc3:3V5 | 5 | 0 |
| *mcd1-1 elg1Δ msh3Δ* | 5 | 5 |
| *mcd1-1 msh3Δ* Smc3:3V5 | 5 | 2 |
| *elg1Δ msh3Δ* Smc3:3V5 | 5 | 3 |
| *mcd1-1 elg1Δ msh3Δ* Smc3:3V5 | 5 | 2 |
| DEAD | 0 | 39 |
| TOTAL | 76 | 72 |
